# A D-amino acid produced by plant-bacteria metabolic crosstalk empowers interspecies competition

**DOI:** 10.1101/2020.06.17.156976

**Authors:** Alena Aliashkevich, Matthew Howell, Gabriella Endre, Eva Kondorosi, Pamela J.B. Brown, Felipe Cava

## Abstract

The bacterial cell wall is made of peptidoglycan (PG), a polymer that is essential for maintenance of cell shape and survival. Many bacteria alter their PG chemistry as a strategy to adapt their cell wall to environmental challenges. Therefore, identifying these factors is important to better understand the interplay between microbes and their habitat. Here we used the soil bacterium *Pseudomonas putida* to uncover cell wall modulators from plant extracts and found canavanine (CAN), a non-proteinogenic amino acid. We demonstrated that cell wall chemical editing by CAN is licensed by *P. putida* BsrP, a broad-spectrum racemase which catalyzes production of D-CAN. Remarkably, D-CAN alters dramatically the PG structure of Rhizobiales (e.g. *Agrobacterium tumefaciens, Sinorhizobium meliloti*), impairing PG synthesis, crosslinkage and cell division. Using *A. tumefaciens* we demonstrated that the detrimental effect of D-CAN is suppressed by a single amino acid substitution in the cell division PG transpeptidase penicillin binding protein 3a. Collectively, this work provides a fascinating example of how interspecies metabolic crosstalk can be a source of novel cell wall regulatory molecules to govern microbial biodiversity.

## Introduction

Bacteria establish a myriad of complex social structures with other living organisms in the biosphere that frequently involve competitive and cooperative behaviours [1, 2]. For instance, many mutualists rely on each other for nutrients and protection [3–6]. Evolution has consolidated these partnerships by selecting specific mechanisms which provide a mutual benefit to the partners, making the interactions more efficient and robust. A representative example of mutualism is the case of legume plants and rhizobia bacteria. Legumes produce flavonoid signals to recruit nitrogen fixing bacteria to the plant. Microbes provide nitrogen in return for energy-containing carbohydrates [7–11]. Ecologists consider that these type of plant-bacteria interactions are more widespread in nature than was previously thought [12, 13].

The development of specific social relationships often requires communication strategies. One such strategy is the production and release of small diffusible molecules, which facilitate interactions between organisms in the distance and often are instrumental to shape the biodiversity, dynamics and ultimately, the biological functions of the ecosystems [14, 15]. Many taxonomically unrelated bacteria produce non-canonical D-amino acids (NCDAAs) to the extracellular milieu in order to regulate diverse cellular processes at a population level. The regulatory properties of NCDAA seem to be specific for each D-amino acid, e.g. D-Met and D-Leu downregulate peptidoglycan (PG) synthesis [16–18], D-Ala represses spore germination [19] and D-Arg affects phosphate uptake [20] (reviewed in [21]).

The modulatory effects of NCDAA on the cell wall require that these molecules replace the canonical D-Alanine located at the terminal position (4^th^ or 5^th^) of the PG peptide stems. NCDAA editing at 4^th^ position is catalysed by LD-transpeptidases (Ldts), which are enzymes involved in PG crosslinking (i.e. dimer synthesis) through the formation of meso-diaminopimelic acid (mDAP-mDAP) peptide bridges [17]. In contrast, incorporation of NCDAA at the 5^th^ is mediated by penicillin binding proteins (PBPs) with DD-transpeptidase activity [22] or by synthesis of modified precursors in the cytoplasmic *de novo* synthetic pathway [17]. Since muropeptides are substrates for many enzymes, PG changes induced by NCDAA can have an obvious impact on the enzymes that synthesize and remodel the PG.

Production of many NCDAAs depends on the enzyme broad-spectrum racemase (Bsr), which converts L-amino acids, protein building blocks, into D-amino acids, regulatory molecules [23]. The wide distribution of Bsr-bacteria [23] and the metabolic investment in producing NCDAA suggests an important physiological role for these molecules. It is worth mentioning that the capacity to incorporate NCDAA in the PG is widespread in bacteria. The fact that non-producer organisms can be also influenced by PG editing suggests that NCDAA can act as engines of biodiversification within poly-microbial communities [20].

Although the implications of NCDAAs in microbial ecology is rapidly growing, yet most studies focus on the production of D-amino acids from their proteinogenic L-counterparts while non-proteinogenic amino acids are much less studied. Here, we report that Bsr of soil bacterium *Pseudomonas putida* (BsrP) can effectively produce D-canavanine (D-CAN) from plant derived L-canavanine (L-CAN), an allelopathic non-proteinogenic amino acid produced by many agronomically important legumes (e.g. alfalfa, jack beans) in high amounts [24–26].

Previous studies have reported that L-CAN causes growth inhibition of non-producer plants due to the induction of systemic protein misfolding associated with the capacity of L-CAN to replace L-Arginine in proteins [27–30]. Our results show that conversion of L-into D-CAN by BsrP eliminates the toxic effect of L-CAN in the growth of *Arabidopsis thaliana*.

Since this is the first time enzymatic D-CAN production is reported we decided to investigate the biological activity of this plant-derived D-amino acid on the physiology of rhizosphere microbes. We found that D-CAN is incorporated in high amounts in the cell wall of certain Rhizobiales species. Cell wall chemical editing by D-CAN affects PG synthesis and structure which causes cell division impairment and fitness loss. Using the plant pathogen *Agrobacterium tumefaciens* we demonstrated that D-CAN deleterious effects on cell wall integrity can be alleviated by just a single amino acid substitution in the cell division PG transpeptidase penicillin binding protein 3a (PBP3a).

## Materials and Methods

### Media and growth conditions

Detailed information about strains and growth conditions is listed in supplementary materials and methods. All strains were grown at the optimal temperature and in LB (Luria Bertani broth) medium unless otherwise stated. Growth of diverse rhizobial species shown in Figure 2 was performed at room temperature.

**Fig. 1.**
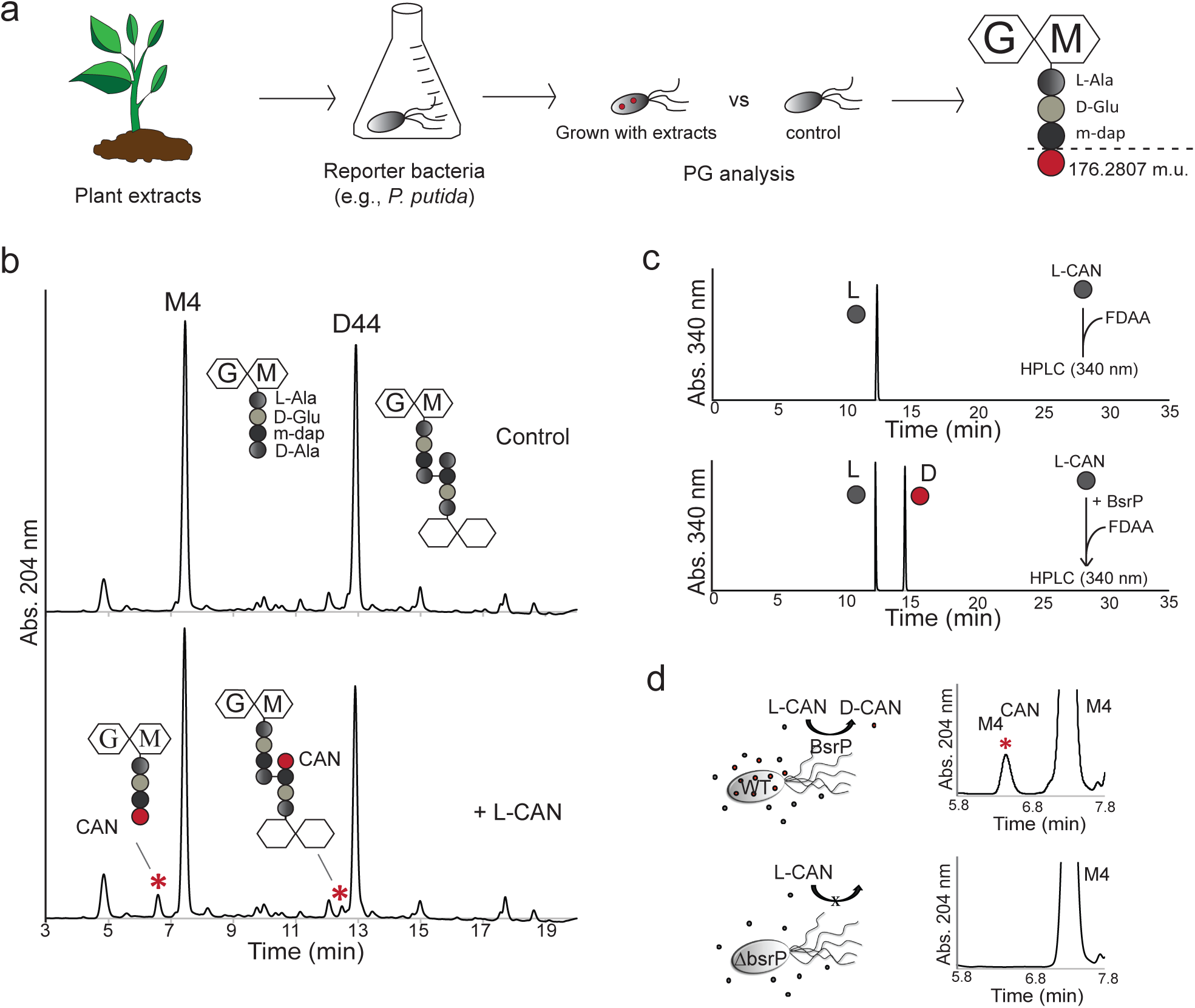
D-canavanine is produced from L-canavanine. (a) Scheme of PG-modifying metabolites identification. Modified M4 was found in the sample grown with *Medicago sativa* (alfalfa) seeds extract. (b) Cell wall analysis of *P. putida*, grown without (control) or with addition of L-canavanine 5 mM. (c) HPLC analysis of Marfey’s derivatized L-canavanine and L-canavanine incubated with *P. putida* broad-spectrum racemase. (d) Cell wall analysis of *P. putida* wt and Δ*bsrP* mutant, grown in the presence of L-canavanine 5 mM.

**Fig. 2.**
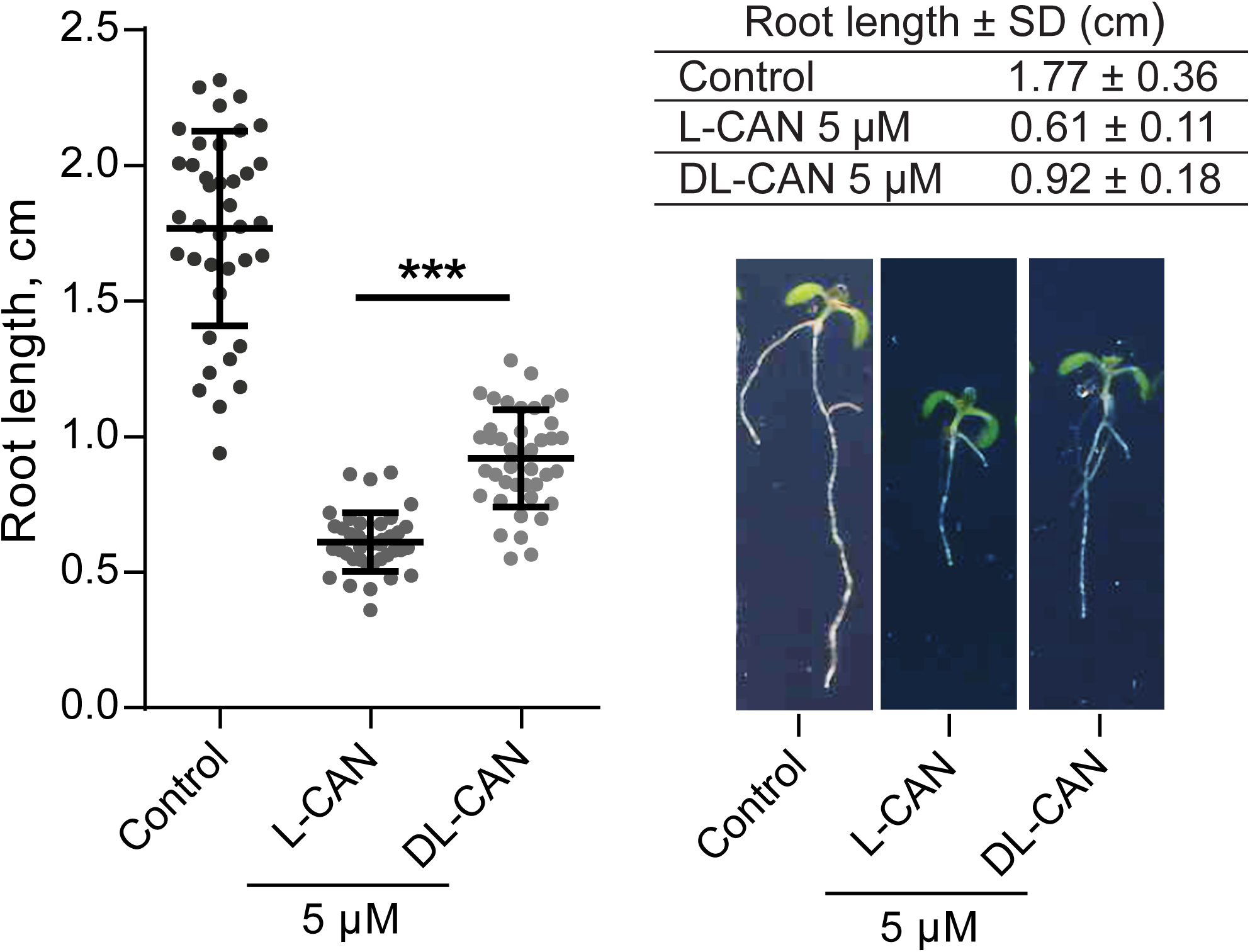
Functionality of D-canavanine is different from L-canavanine. Root length in *A. thaliana* grown on ½ Murashige-Skoog agar supplemented with L- or DL-canavanine 5 µM or not (control). Pictures show representative plants. P value < 0.0001 (***).

### Seed extract preparation and use of *P. putida* as a reporter

3 gr of seeds (e.g. *Medicago sativa*) were mashed and soaked in 10 mL of water overnight followed by centrifugation at 5,000 rpm to remove the particulate fraction. The supernatant was next (i.e. extract) filter-sterilized and concentrated 5x. *P. putida* were grown either in LB medium or in LB medium supplemented with seed extract to a final concentration 1x. Cultures were grown up to stationary phase prior PG purification and analysis by liquid chromatography and by mass spectrometry.

### Peptidoglycan analysis

PG isolation and analysis were done according previously described methods [31, 32]. In brief, PG sacculi were obtained by boiling bacterial cells in SDS 5%. SDS was removed by ultracentrifugation, and the insoluble material was further digested with muramidase (Cellosyl). Soluble muropeptides were separated by liquid chromatography (high-performance liquid chromatography and/or ultra high-pressure liquid chromatography) and identified by mass spectrometry. A detailed protocol is described in supplementary materials and methods.

### Protein expression and purification

*P. putida* gene PP3722 encoding broad-spectrum racemase was amplified with FCP1097 (5’-AAAACATATGCCCTTTCGCCGTACC-3’) and FCP1098 (5’-AAAAGCGGCCGCGTCGACGAGTAT-3’) primers and cloned in pET22b for expression in *E. coli* Rosetta 2 (DE3) cells, resulting in C-terminal His-tagged protein.

Protein was purified using Ni-NTA agarose column (Qiagen). A detailed protocol is described in supplementary materials and methods.

### Racemase activity assay

5 µg of purified racemase and various concentration of L-canavanine in 50 µl of 50 mM sodium phosphate buffer pH 7.5 were incubated at 37 °C for 30 min, then heat inactivated (5 min, 100°C), and centrifuged (15,000 rpm, 10 min). Supernatant was derivatized with Marfey’s reagent [33] and resolved by high-performance liquid chromatography as described previously [23]. Detailed protocols are available in supplementary materials and methods.

### *BsrP* mutant construction in *P. putida*

For deletion of PP3722 in *P. putida* the upstream and downstream regions of the gene were amplified from purified genomic DNA with primers FCP1145 (5’-AAAATCTAGATCATCAGCAGCGACAT-3’) and FCP1092 (5’-CAATGGCAATTGGTGATTACTCGTGTTC-3’); FCP1093 (5’-GAGTAATCACCAATTGCCATTGAAAGGAG-3’) and FP1146 (5’-AAAATCTAGAGCGACGTCACGC-3’) respectively. The upstream and downstream fragments were combined with FCP1145 and FCP1146 into a 1010 bp fragment, and inserted into pCVD442 [34]. *E. coli* DH5α λPIR was used in the cloning and the resulting plasmid pCVD442*bsrP* was confirmed by sequencing. In-frame deletion was introduced by allele replacement via homologous recombination. In short, exconjugants were obtained by conjugating with Sm10 λPIR containing pCVD442*bsrP* and selected on LB plates with chloramphenicol 25 μg/ml and carbenicillin 1,000 μg/ml. Exconjugants were grown in LB with 10% sucrose (w/v) medium overnight and then plated on LB plates with chloramphenicol 25 μg/ml and 10% (w/v) sucrose. Colonies sensitive to carbenicillin were confirmed by PCR.

### *A. thaliana* growth

*A. thaliana* was grown in ½ MS agar medium (half strength of Murashige and Skoog basal salt mixture (Sigma), 0.5% sucrose, 1% agar, with pH adjusted to 5.7) with or without canavanine supplementation. Ethanol sterilized seeds were pre-incubated on the plates in the darkness at 4°C for 3 days before moving to the *in vitro* chamber with day/night cycle 16/8 hours, 22°C/18°C. Root length was measured after 10 days of growth in the chamber with Fiji [35]. Pictures of the root hairs were taken with stereomicroscope Nikon SMZ1500 (Tokyo, Japan).

### Growth curves and relative growth

At least three replicates per strain and growth condition were grown in 200 μl of LB alone or supplemented with canavanine in a 96-well plate at 30°C with 140 rpm shaking in a BioTek Eon Microplate Spectrophotometer (BioTek, Winooski, VT, USA). The A600 was measured at 10 minutes intervals. Relative growth was calculated as a percentage of growth in the presence of DL-canavanine compared to growth without canavanine.

### Phase contrast microscopy

Stationary phase bacteria were placed on 1% agarose LB pads. Phase contrast microscopy was done using a Zeiss Axio Imager.Z2 microscope (Zeiss, Oberkochen, Germany) equipped with a Plan-Apochromat 63X phase contrast objective lens and an ORCA-Flash 4.0 LT digital CMOS camera (Hamamatsu Photonics, Shizuoka, Japan), using the Zeiss Zen Blue software.

### Quantification of cell constrictions

Exponentially growing cells (OD_600_=0.4-0.6) in ATGN medium [36] were imaged on 1% agarose ATGN pads using phase contrast microscopy (inverted Nikon Eclipse TiE (Tokyo, Japan) with a QImaging Rolera em-c2 1K EMCCD camera (Surrey, British Columbia. Canada), and Nikon Elements Imaging Software) as described previously [37]. Cell length and constrictions were detected using MicrobeJ software [35]. Old poles were identified as having a larger maximum width compared to the new poles. The longitudinal position of cell constrictions was then plotted against cell length. A longitudinal position of 0 represents the true midcell while positive values approach the new pole and negative values approach the old cell.

### Suppressor mutants

To obtain suppressor mutants, *A. tumefaciens* was grown at optimal conditions overnight (see supplementary methods), and serial dilutions were inoculated on the LB plates containing DL-CAN 10 mM. Plates were incubated at room temperature until suppressor mutant colonies arose. For confirmation of the resistance, the selected colonies were passed through LB plates before being tested on LB plates containing DL-CAN 10 mM.

### Whole-genome sequencing and single-nucleotide polymorphism analysis

Genomic DNA was isolated from suppressor mutants and the parental strain of *A. tumefaciens*. Indexed paired-end libraries were prepared and sequenced in a MiSeq sequencer (Illumina, San Diego, CA, USA) according to the manufacturer’s instructions.

Data quality control was performed with FastQC v0.11.5 [38] and MultiQC v1.5 [39]. The raw data in FASTQ format was trimmed using Trimmomatic v0.36 with arguments ‘ILLUMINACLIP:adapters.fa:2:30:10’,’SLIDINGWINDOW:5:30’ and ‘MINLEN:50’ [40]. The exact adapter sequences that were used can be retrieved from the supplementary materials and methods. The trimmed FASTQ was aligned to genome GCF_000092025.1_ASM9202v1 (A. tumefaciens, [41]) using the ‘mem’ algorithm in BWA v0.7.15-r1140 [42] with default parameters and subsequently converted to sorted BAM format. Optical duplicates were marked using picard tools v2.18.2 with default arguments [43]. Finally, variants were called in freebayes v1.1.0-dirty using the parameters ‘-p 1’, ‘--min-coverage 5’ and ‘--max-coverage 500’ [44].

### Reconstruction of suppressor mutant *pbp3a*^K537R^ in *A. tumefaciens*

For reconstruction of point mutation in *pbp3a*^K537R^ in *A. tumefaciens*, a 650 bp fragment containing the mutated nucleotide was amplified from purified genomic DNA with primers FCP3354 (5’-AAAAGGATCCCGACACCGTTGG-3’) and FCP3355 (5’-AAAAGGATCCATAAGACACGAGCA-3’) and inserted into pNPTS139 plasmid [45]. *E. coli* DH5α λPIR was used in the cloning and the resulting plasmid pNPTS139*pbp3a*^K537R^ was confirmed by sequencing.

Nucleotide substitution in *A. tumefaciens pbp3a* gene (atu2100) was done according to an established allelic-replacement protocol [46]. In short, exconjugants were obtained by conjugating with *E. coli* S17-1 λPIR containing pNPTS139*pbp3a*^K537R^ and selected on ATGN plates with kanamycin 300 μg/ml. Exconjugants were grown in ATGN medium overnight and then plated on ATSN plates with 5% (w/v) sucrose [36]. Colonies sensitive to kanamycin were streak-purified twice on ATSN plates and sequenced.

### PBP3a protein folding prediction

Prediction of PBP3a protein was done by Phyre2 [47].

## Results

### Bacterial racemization of canavanine licenses its incorporation into the cell wall

To identify new environmental modulators of the bacterial cell wall we tested the capacity of diverse plant extracts to induce changes in the PG chemical structure of soil associated bacteria. We found that our reporter strain *Pseudomonas putida* displayed new muropeptides when we supplemented its growth medium with alfalfa (*M. sativa*) seed extract. By mass spectrometry, we identified that the modification corresponded to a molecule of 176.2807 mass units that was replacing the D-Alanine normally found at fourth position of the peptides stems within the bacterial PG (Fig. 1a). *In silico* analyses suggested L-canavanine (L-CAN), a non-proteinogenic amino acid similar to L-arginine and found in legumes as the most likely candidate. Consistently, supplementation of *P. putida* with pure L-CAN produced the same monomeric muropeptide, now renamed as M4^CAN^, but also its crosslinked dimeric form D44^CAN^ (Fig. 1b). Since the fourth position in the peptide moiety of muropeptides is normally restricted to D-amino acids, we hypothesized that *P. putida* might have produced D-CAN from L-CAN. In fact, we found that *P. putida* genome encodes a putative broad-spectrum racemase orthologue (PP3722). To test whether PP3722 could racemize canavanine we purified the protein and performed *in vitro* racemization (reversible interconversion between L-AA and D-AA enantiomers) assays using pure L-CAN as substrate. Indeed, using High Performance Liquid Chromatography (HPLC) we observed that PP3722 converted L-CAN into D-CAN and hence we named this protein as BsrP for <u>B</u>road-<u>s</u>pectrum <u>r</u>acemase in *<u>P</u>. putida* (Fig. 1c).

Consistently, deletion of *bsrP* in *P. putida* produced a strain incapable to make D-CAN-containing muropeptides in L-CAN supplemented cultures (Fig. 1d). *P. putida* Δ*bsrP* was only able to produce a PG edited with CAN when this was exogenously added as D-form. Since we did not succeed in purifying D-CAN, we used DL-CAN racemic mixture as a source of D-CAN (DL-CAN) (Fig. S1a). In agreement, D-CAN containing supernatants (from wild-type (wt) *P. putida*) induced production of D-CAN muropeptides in *E.coli*, a bacterium that lacks broad-spectrum racemase (Fig. S1b), further supporting that PG modification by D-CAN is Bsr-independent. As expected, no D-CAN muropeptides were induced in *E. coli* when this bacterium was cultured with preconditioned media from the Δ*bsrP* strain. Collectively, these results indicate that bacterial broad-spectrum racemase BsrP can change the chirality of plant-derived amino acid L-CAN, thereby licensing its D-form for PG editing.

### Enantiomerization changes the functionality of canavanine

Previous studies showed that production of L-CAN by legumes underlies a defensive strategy against certain competitors (e.g. plants, insects) [27, 48, 49] based on the incorporation of this toxic atypical amino acid into proteins due to its chemical similarities with L-arginine [28–30]. Compared to L-CAN, there is virtually no information about D-CAN. Thus, to understand the biological role of this D-amino acid we first checked if D-CAN displayed the same activity as L-CAN. In agreement with previous reports, L-CAN inhibited root growth of *A. thaliana* seedlings at 5 µM concentration with the resulting root length almost 3 times shorter than in control (Fig. 2). However, the average root length in the presence of DL-CAN 5 µM was 1.5 times longer than that grown with the same concentration of L-CAN suggesting that CAN enantiomers have different functions. Indeed, additional experiments comparing root lengths at L-CAN 5 µM versus DL-CAN 10 µM (i.e. 5 µM D-CAN + 5 µM L-CAN), and L-CAN 10 µM versus DL-CAN 20 µM (i.e. 10 µM D-CAN + 10 µM L-CAN), where in both cases amount of L-form is the same, revealed no significant differences between them (Fig. S2a) and suggests that only L-CAN inhibits root development in *A. thaliana*. Interestingly, in addition to tap root length, development of lateral roots and root hairs were also affected by L-CAN, but not by D-CAN (Fig. S2b). Collectively, these results stress the idea that CAN enantiomers have different activities.

### D-CAN severely alters cell wall composition and abundance in Rhizobiales

To ascertain the physiological role of D-CAN we investigated its effect on bacterial growth using diverse bacteria species that can potentially be exposed to this D-amino acid in the natural environment. We found that Rhizobiales were the most affected species by DL-CAN (Fig. 3a) while *P. putida* growth was not affected even at high levels of DL-CAN (up to 10 mM) (Fig. S3) suggesting that producer species (i.e. encoding a broad-spectrum racemase) might have developed tolerance to D-CAN.

**Fig. 3.**
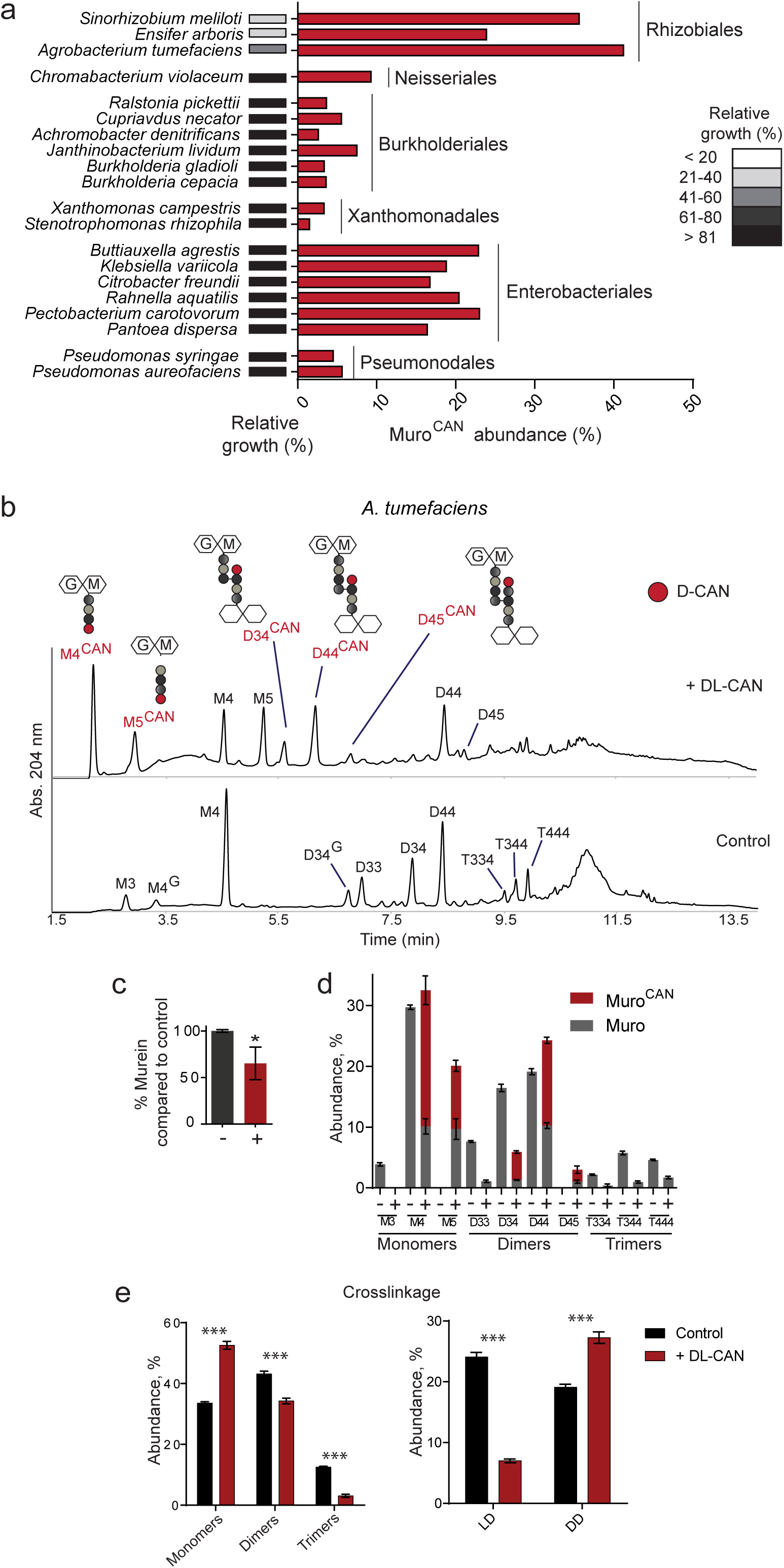
High D-canavanine incorporation changes structure and amount of peptidoglycan in *A. tumefaciens*. (a) Sensitivity of soil and ubiquitous bacteria to DL-canavanine. Relative growth was calculated for bacteria grown in the presence of 5 mM DL-canavanine. D-canavanine incorporation was measured for bacteria supplemented with 2.5 mM DL-canavanine. (b) Representative PG profiles of *A. tumefaciens* supplemented with DL-canavanine 10 mM or not (control). Illustrations show D-canavanine-containing muropeptides. (c) PG amount quantification in 10 mM DL-canavanine supplemented *A. tumefaciens* cultures normalized to control (no canavanine). P-value < 0.05 (*). (d) Abundance of D-canavanine-containing muropeptides in *A. tumefaciens* supplemented with 10 mM DL-canavanine. Monomer M4^G^ and dimer D34^G^ are calculated as part of non-modified M4 and D34. (e) Abundance of monomers, dimers and trimers in *A. tumefaciens* supplemented with 10 mM DL-canavanine. Abundance of LD- and DD-crosslinked muropeptides in *A. tumefaciens* supplemented with 10 mM DL-canavanine. P value < 0.0001 (***).

Although D-CAN induced PG modifications in all species tested, Rhizobiales displayed the highest levels of muro^CAN^, i.e. ca. 40% of the muropeptides were edited by D-CAN both in the 4^th^ and 5^th^ positions of the peptide moieties (Fig. 3a, b, Fig. S4a). Therefore, we hypothesized that D-CAN might be interfering in cell wall biosynthesis, in a similar way as has been reported for other NCDAAs (e.g. D-Met [20, 50]). Indeed, *A. tumefaciens* cells treated with DL-CAN contained less PG than non-treated cells (Fig. 3c) or cells treated with L-CAN (Fig. S4b). To investigate the consequences of D-CAN incorporation on the PG architecture, we added increasing concentrations of DL-CAN to *A. tumefaciens* and monitored fluctuation of the different PG components. Our results show that D-CAN causes a dramatic increase in pentapeptides (M5 and D45) (Fig. 3b, d), and a reduction in crosslinkage due to lower amount of LD-crosslinked muropeptides (Fig. 3e). L-CAN alone did not change *A. tumefaciens* PG crosslinkage at tested concentration (Fig. S4c).

To know if the effects of D-CAN in *A. tumefaciens*’ PG extend to other Rhizobiales, we analyzed both PG composition and amount in the legume symbiont *Sinorhizobium meliloti*. As in *A. tumefaciens*, we found the same types of D-CAN modified muropeptides, reduction in PG density and crosslinkage in *S. meliloti* treated with D-CAN (Fig. S5a, S5b, S5c). Interestingly, we had to use lower concentration of the compound, since *S. meliloti* was more sensitive to D-CAN than *A. tumefaciens*. These results suggest that D-CAN downregulates PG synthesis and crosslinkage likely through its incorporation in the cell wall.

Given the effects of D-CAN in *S. meliloti*, we decided to explore the effect of this D-amino acid on *Medicago sativa*, a legume which produces L-CAN and establishes symbiosis with *Sinorhizonium medicae* for nitrogen fixation. Pre-treatment of *S. medicae* with DL-CAN delayed nodulation, reduced the nodule number and caused early senescence and disintegration of the nitrogen-fixing nodule zone (Fig. S6a, b). As a consequence of the lack of active persistent nitrogen-fixing cells, the aerial part of plants was underdeveloped and similar to the non-infected, nitrogen-starving plants (Fig. S6c). Collectively, our data demonstrates that D-CAN activity can affect the fitness of certain rhizobia and as a consequence, their symbiotic relationship with plants.

### D-CAN impairs viability and cell separation

To gain further insights on D-CAN’s mechanism of action we cultured *A. tumefaciens* with or without L- or DL-CAN and monitored growth and morphology. Our results showed that D-CAN inhibited growth of *A. tumefaciens* in liquid culture and induced lysis, branching and bulging (Fig. 4a). No significant changes in growth or morphology were caused by L-CAN (Fig. 4a) further strengthening the idea that these enantiomers have different functions. To get more quantitative insights of the morphological defects caused by D-CAN we measured cell length, longitudinal position of the constriction (Fig. 4b), and the number of constrictions per cell (Fig. 4c). While in the untreated culture, or in cultures treated with L-CAN, *A. tumefaciens* division sites localized slightly closer to the new pole (Fig. 4b, Fig. S7a), in DL-CAN treated cultures cells were up to 1.5 times longer and the position of the constrictions exhibited a more scattered pattern (Fig. 4b). In addition, untreated cells and cells treated with L-CAN had 0 or 1 constriction per cell, while DL-CAN induced up to 3 constrictions per cell (Fig. 4c, Fig. S7a). As before, *S. meliloti* grown on DL-CAN recapitulated the results obtained with *A. tumefaciens* on growth, morphology and number of constrictions (Fig. S7b, c, d) further supporting that D-CAN interferes with the cell division.

**Fig. 4.**
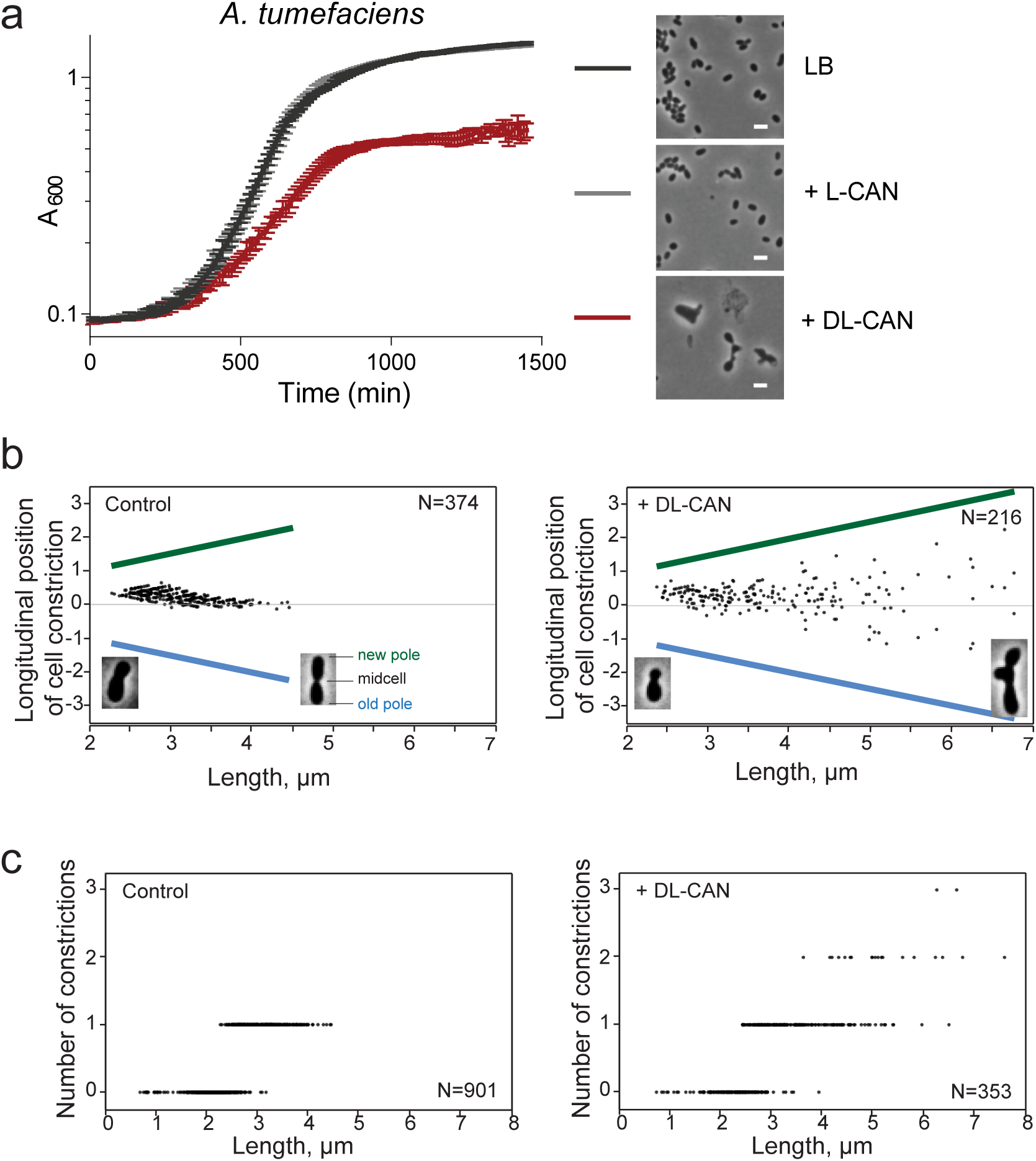
D-canavanine inhibits growth of *A. tumefaciens* and leads to aberrant cell morphology. (a) Growth curves of *A. tumefaciens* in the absence (control) or presence of L- or DL-canavanine 10 mM, and phase contrast images of *A. tumefaciens* cells without (control) or supplemented with L- or DL-canavanine 10 mM. Scale bar 2 µm. (b) Longitudinal position of cell constriction in *A. tumefaciens* cells without (control) or with DL-canavanine 10 mM. New pole is marked by green color, old pole – by blue. (c) Number of constrictions per cell in *A. tumefaciens* grown without (control) or with DL-canavanine 10 mM.

### D-CAN interfere with a cell division transpeptidation

To identify the molecular targets of D-CAN, we screened for suppressor mutants resistant to DL-CAN. Characterization of the single-nucleotide polymorphism by genome sequencing revealed a K537R substitution in the primary cell division transpeptidase PBP3a (*atu2100*) [51, 52]. Phyre2 alignments [47] of *A. tumefaciens* PBP3a to crystallized PBP3 proteins localized K537 in the loop between β5 and λ11, close to the active-site cleft (Fig. 5a).

**Fig. 5.**
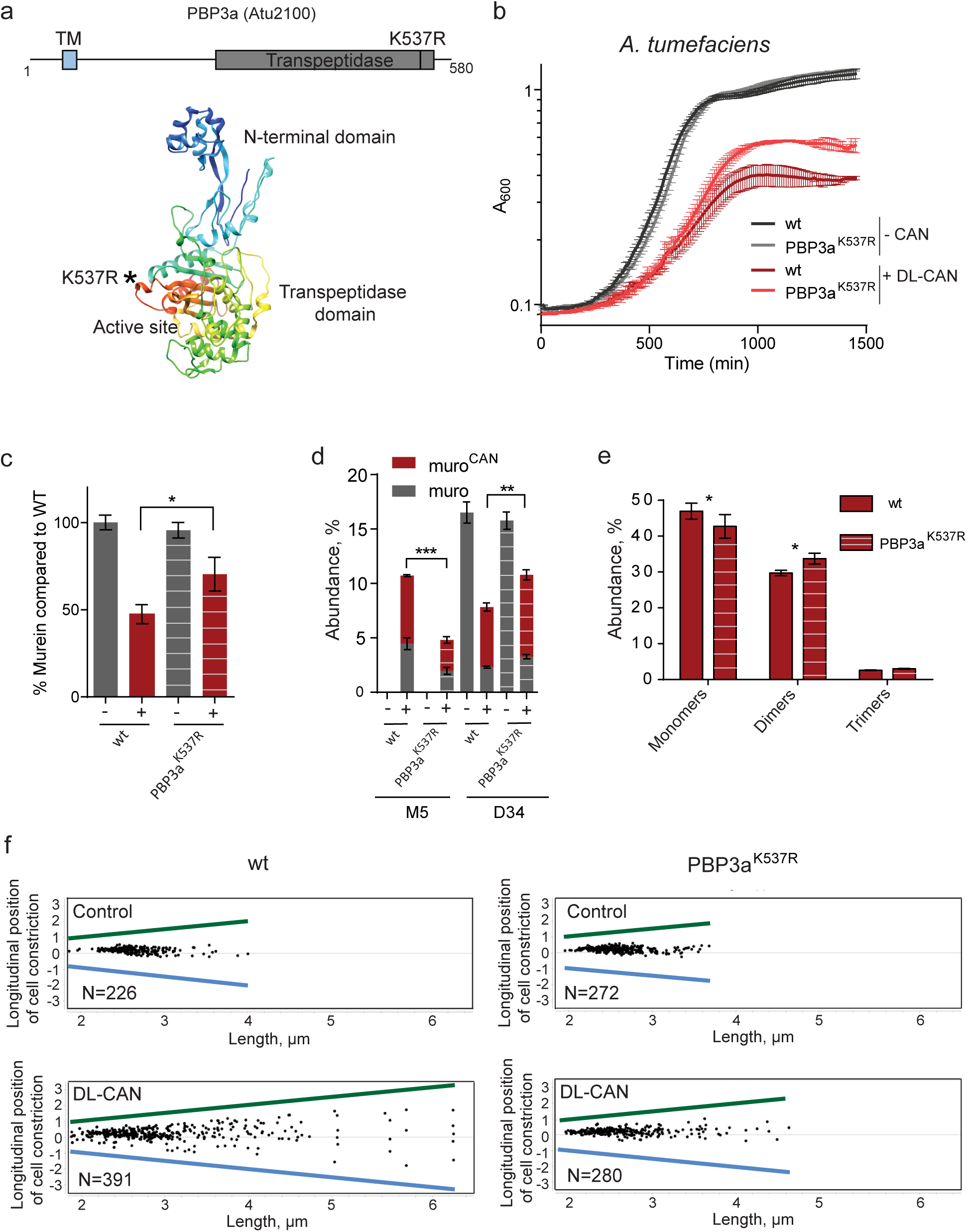
K537R amino acid change in *A. tumefaciens* PBP3a protein provides resistance to D-canavanine. (a) Position of the PBP3a K537R amino acid change in the protein scheme and in the protein structural prediction. (b) Growth curves of *A. tumefaciens* wild-type and PBP3a^K537R^ in the presence of DL-canavanine 10 mM. (c) PG amount quantification in 10 mM DL-canavanine supplemented *A. tumefaciens* wild-type and PBP3a^K537R^ cultures normalized to wild-type control (no canavanine). P-value < 0.05 (*). (d) Quantification of the monomer (M5) and dimer (D34) abundance in *A. tumefaciens* wild-type and PBP3a^K537R^ grown with DL-canavanine 10 mM. P-value < 0.005 (**) and < 0.0001 (***). (e) Abundance of monomers, dimers and trimers in *A. tumefaciens* wild-type and PBP3a^K537R^ supplemented with 10 mM DL-canavanine. P-value < 0.05 (*). (f) Longitudinal position of cell constriction in *A. tumefaciens* wild-type and PBP3a^K537R^ cells without (control) or with DL-canavanine 7.5 mM. New pole is marked by green color, old pole – by blue.

Reconstruction of the K537R mutation (i.e. *A. tumefaciens* PBP3a^K537R^) recapitulated the suppressor tolerance to DL-CAN (Fig. 5b). Interestingly, K537R substitution appeared to be specific since it did not suppress the growth inhibitory effect of D-amino acids other than D-Arg, a chemical analogue of D-CAN (Fig S8). No difference in the growth of the wt and PBP3a^K537R^ strains was detected in the absence or presence of L-CAN (Fig. S9a). In addition to growth, PG reduction was partially alleviated in the PBP3a^K537R^ strain (Fig. 5c). Both wild-type vs the PBP3a^K537R^ strains showed similar levels of D-CAN containing muropeptides (muro^CAN^) in cultures supplemented with DL-CAN indicating that the suppressing role of the PBP3a^K537R^ mutations is not associated with a reduction of D-CAN incorporation in the PG (Fig. S9e).

Consistent with the idea that D-CAN inhibits PBP3a activity, the PBP3a^K537R^ strain showed a reduction in the accumulation of pentapeptides (i.e. M5) compared to that of the wild-type in the presence of D-CAN (Fig. 5d, Fig. S9b). Overall crosslinkage levels and particularly LD-crosslinkage also improved in the PBP3a^K537R^ strain (Fig. 5e, Fig. S9c), while no difference between strains was observed in control condition (Fig. S9d). Similarly, altered cell length and constriction positioning in the presence of DL-CAN improved in the PBP3a^K537R^ strain compared to wt (Fig. 5f), while no difference was observed in the control condition or in the presence of L-CAN (Fig. 5f, Fig. S9f). Collectively, these data suggest that D-CAN interfere with PG transpeptidation at cell division.

## Discussion

Bacteria can edit the canonical chemistry of their cell wall as a strategy to cope with environmental challenges [53–55]. As PG can be modified by secreted molecules, we reasoned that we could use bacteria as a biochemical trap to discover elusive environmental modulators of the cell wall. To test this, we exposed plant-derived soluble extracts to the soil bacteria *P. putida* and discovered canavanine (CAN) as a new PG modulator. The fact that L-CAN was previously reported to be produced by legume plants [24–26] further supported the efficacy of our screening. However, CAN was found at the terminal position of the PG peptide moieties, which is reserved for D-amino acids [16]. Remarkably, we found that *P. putida* encodes a broad-spectrum racemase (Bsr) that changes the chirality of CAN to permit its incorporation in the bacterial PG. Collectively, these observations underscore a fascinating example of interspecies metabolic crosstalk where a plant-derived metabolite (L-CAN) is transformed by a bacterial enzyme (BsrP) into a previously unrecognized molecule (D-CAN) (Figure 6). Discovery of D-CAN adds to a growing list of metabolites produced as a result of plant-soil feedbacks and contributes to chemical ecology. [56–58].

**Fig. 6.**
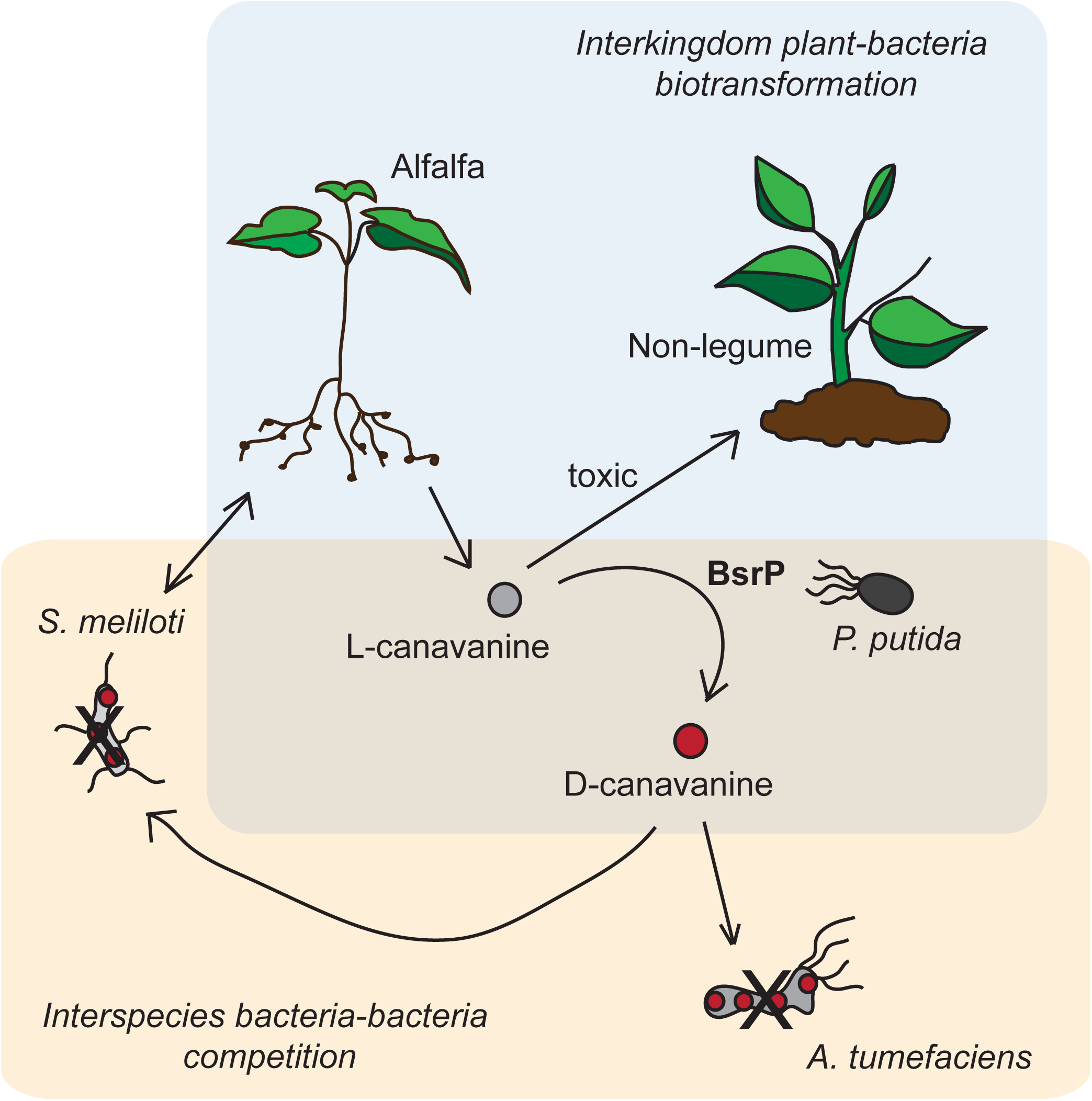
Model illustrating the impact of L- to D-CAN conversion on the soil-plant ecosystem. CAN enantiomerization by Bsr bacteria (e. g. *P. putida*) detoxifies L-CAN for non-legume plants. D-CAN inhibits Rhizobiales bacteria (e. g. *S. meliloti, A. tumefaciens*), thus modulating microbial diversity in the soil.

Since amino acid enantiomers have different functions, racemization of CAN may lead to multiple environmental effects. On one side, Bsr racemization of L-CAN to D-CAN decreases the concentration of the L-CAN, alleviating its toxic effect on plants [27]. In addition, Bsr produces D-CAN, a molecule that alters bacterial PG composition (Figure 6).

PG editing is a mechanism by which the environment can regulate the cell wall structure and biosynthesis. Whether this regulation is positive or detrimental seems to depend both on the type of D-amino acid and on the bacteria species. For instance, although *Vibrio cholerae* produces and incorporates both D-Arg and D-Met in its PG, only the latter has an effect on cell wall synthesis [20]. In the particular case of D-CAN, it seems clear that the most sensitive species were those with polar growth and higher levels of D-CAN in the PG. Indeed, many Rhizobiales elongate unidirectionally by adding PG to the new pole, generated after cell division [59]. When new cell compartment gets bigger in length and width, the zone of active PG growth together with division proteins localize to midcell prior to cell division. *A. tumefaciens* encodes multiple LD-transpeptidases (e.g. 14 Ldts in *A. tumefaciens* compared to just two predicted orthologues in *P. putida*) and different Ldts are localized to the new pole or midcell, and presumably important for both polar growth and division [51]. Ldts are the enzymes that perform mDAP-mDAP crosslinks, which are very abundant in Rhizobiales (40-50% in *A. tumefaciens*) compared to e.g. *P. putida* (ca. 1 %), and catalyze PG editing in the 4^th^ position of the peptide moieties [17]. Therefore, free D-CAN might act as a competitive substrate on Ldts to prevent their LD-crosslinking activity in favor of high D-CAN incorporation. In fact, D33 and D34 LD-crosslinked dimers are significantly reduced in the present of D-CAN. The high number of Ldt paralogs in these species suggest they are important for the lifestyle of these organisms and thus might be difficult to assess whether a D-CAN deleterious effect can be suppressed in a Ldt-deficient strain. Another target of D-CAN inhibition might be DD-carboxypeptidases, enzymes that remove the terminal D-Ala from pentapeptides (M5). Accumulation of both the canonical (D-Ala-terminated pentapeptides) and the non-canonical (D-CAN-pentapeptides) in the presence of D-CAN strongly suggest that free D-CAN decreases the activity of *A. tumefaciens* DD-carboxypeptidases.

Interestingly, our suppressor analyses did not identify any mutations in Ldts or DD-carboxypeptidases that improved the growth of *A. tumefaciens* in the presence of D-CAN. The high number of Ldt and DD-carboxypeptidase paralogues (14 and 4 predicted, respectively) makes very unlikely that a single mutation in these proteins would show a suppressor effect. Instead, we discovered that a K537R point mutation in the PBP3a (*atu2100*) is sufficient to alleviate D-CAN sensitivity in *A. tumefaciens.* There are two important evidences in agreement with the idea of D-CAN targeting PBP3a: i) PBP3a has been reported to localize at the septum and be involved in cell division. Consistently, D-CAN induces branching and bulging in the wt and the PBP3a K537R mutation suppresses this phenotype. Ii) PBP3a is a DD-transpeptidase. Inhibition of these enzymes reduce crosslinkage levels and increase accumulation of the monomeric substrates (pentapeptide and/or tetrapeptide monomers, i.e. M5 and M4, respectively). Indeed, D-CAN induces M5 accumulation in the wt, which is suppressed in the K537R mutant. Overall DD-crosslinkage is not reduced by D-CAN, but it’s possible that D-CAN targets PBP3a and other PBPs are not inhibited.

The nature of the observed increase in D34 dimers in the K537R mutant seems to be indirect while yet connected to the presence of D-CAN. D34 dimers are formed between two monomer tetrapeptides (M4) by LD-transpeptidases, not by PBP3a, which is DD-transpeptidase and would produce a D43 dimer instead. One might speculate that PG analysis gives overview on overall PG structure, however structural changes in the septal PBP3a might have allosteric consequences on nearby enzymes within a same protein complex. In this line, it has been reported that several Ldt enzymes predominantly localize to the midcell at cell division in *A. tumefaciens* [51]. Therefore, it might possible that PBP3a K537R mutation influences the activity of septal Ldts. Alternatively, PBP3a K537R mutation might induce allosteric regulatory changes in DD-carboxypeptidase at the septum, leading to local consumption of pentapeptides at cell division and increase in the levels of M4, which as Ldt substrates, can boost formation of D34.

Collectively, these results suggest that D-CAN incorporation downregulates PBP3a, among other cell wall associated activities, to inhibit PG synthesis, cell division and induce cell lysis (Figure 6). We hypothesize that K537R substitution might change the properties of the loop between β5 and λ11, which is proximal to the active-site cleft to preserve PBP3a activity while making it insensitive to D-CAN. Understanding the structural changes that K-R mutation induces in the PBP3a structure might provide insights about the underlying mechanisms behind D-CAN tolerance in other bacterial species.

Finally, we have demonstrated that D-CAN affects *S. medicae’s* capacity to facilitate nitrogen-fixation to *M. sativa* (Figure 6). Whether this phenomenon occurs as a consequence of D-CAN impairing the symbiont’s general fitness or a more specific cellular process is something that still needs to be determined. However, recent studies have shown that a DD-carboxypeptidase is critical for bacteroid (specialized nitrogen-fixing cells) differentiation in *Bradyrhizobium* spp. [60, 61], which is consistent with our results of D-CAN downregulating these PG enzymes.

All in all, the ubiquity of bacteria encoding Bsr enzymes strongly suggests that amino acid racemization is an evolutionary driver of cell wall chemical plasticity in the environment. Future research on these enzymes will uncover more interkingdom/interspecies regulatory networks as well as shed new light on how the chirality of amino acids can impact the biodiversity in natural ecosystems.

## Acknowledgements

We thank all the members of the Cava lab for helpful discussions. Research in the Cava lab is supported by The Swedish Research Council (VR), The Knut and Alice Wallenberg Foundation (KAW), The Laboratory of Molecular Infection Medicine Sweden (MIMS) and The Kempe Foundation. Research in the Kondorosi lab is supported by the Frontline Research project KKP129924 from the Hungarian National Office for Research, Development and Innovation and by the Balzan research grant to É. Kondorosi. Research in the Brown lab is supported by the National Science Foundation, IOS1557806.

