## supplementary text and figures for "A D-amino acid produced by plant-bacteria metabolic crosstalk empowers interspecies competition"

Running title: Interspecies competition by D-amino acids.

Keywords: D-amino acids, canavanine, peptidoglycan, cell division, *Agrobacterium tumefaciens*, Rhizobiales, PBP3a.

### Supplementary materials and methods

#### Racemase activity assay

Supernatant of racemization reaction was derivatized with Marfey's reagent (L-FDAA, 1-fluoro-2-4-dinitrophenyl-5-L-alanine amide) [1]. First, an equal volume of $NaHCO_3$ 0.5 M was added to the sample; then, 6 µl of this reaction were reacted with L-FDAA (10 mg/ml in acetone) at 80 ºC for 3 min. HCl 2N was used to stop the reaction and the samples were filtered. The products were separated with a linear gradient of triethylamine phosphate/acetonitrile in HPLC with an Aeris peptide column (250 x 4.6 mm; 3.6 µm particle size) (Phenomenex) and detected at 340 nm. Unracemized L-canavanine was used as control.

#### *E. coli* grown in preconditioned medium

Preconditioned medium (PCM) was obtained from *P. putida* wild-type grown in LB with or without L-CAN until stationary phase. *P. putida* cells were removed from PCM by centrifuging and then filtering supernatant (0.22 µm filters). Stationary phase pre-cultures of *E. coli* were diluted 1/1000 in PCM, fresh 10X LB was added to final concentration of 1X. *E. coli* was grown until stationary phase (about 16 hours). PG was extracted and analyzed.

#### Peptidoglycan isolation

Bacterial cells were harvested by centrifugation (4,000 rpm, 20 min) and boiled in SDS 5% (w/v) for 2 h. Peptidoglycan was pelleted by centrifuging for 13 min at 60,000 rpm at 20 °C (TLA100.3 Beckman rotor; OptimaTM Max ultracentrifuge Beckman, Beckman Coulter, California, USA). Pellets were washed 3-4 times by repeated cycles of centrifugation and resuspension in water.

Samples of *P. putida* were additionally treated with Pronase E (100 μg/ml, 1 h, 60°C) for Braun's lipoprotein removal. The reaction was heat-inactivated, and sacculi were further washed by ultracentrifugation as described above.

The pellet from the last washing was digested with muramidase (Cellosyl 100 μg/ml) for 16 h at 37°C. Muramidase digestion was heat inactivated. Coagulated protein was removed by centrifugation for 15 min at 15,000 rpm. For sample reduction, samples were first adjusted to pH 8.5–9.0 with borate buffer, and then a freshly prepared $NaBH_4$ 2 M solution was added to a final concentration of 10 mg/ml. After 20 min at room temperature, samples were adjusted to pH 3.5 with phosphoric acid and filtered (0.2 μm filters).

**Peptidoglycan analysis**

Analysis of muropeptides was performed on an ACQUITY Ultra Performance Liquid Chromatography (UPLC) BEH C18 column, 130Å, 1.7 μm, 2.1 mm x 150 mm (Waters Corporation, USA) and detected at Abs. 204 nm with ACQUITY UPLC UV-visible detector. Muropeptides were separated at 35°C with inorganic buffers using a linear gradient from buffer A (sodium phosphate buffer 50 mM pH 4.35) to buffer B (sodium phosphate buffer 50 mM pH 4.95 methanol 15% (v/v)) with a flow of 0.25 mL/min in a 20 minutes run. Muropeptides were separated with organic buffers at 45°C using a linear gradient from buffer A (formic acid 0.1% (v/v) in water) to buffer B (formic acid 0.1% (v/v) in acetonitrile) in a 18 minutes run with a 0.25 ml/min flow.

Relative abundance of individual muropeptides was quantified from the relative area of the corresponding peak compared to the total area of the chromatogram. Muropeptides' abundance was statistically compared using unpaired t-test.

Relative total PG amounts were calculated by quantification of total areas (total intensities) of the chromatograms from three biological replicates normalized to the same $OD_{600}$/mL.

**MS analysis**

Samples were analyzed by UPLC chromatography coupled to MS/MS analysis, using a Xevo G2-XS QTof system (Waters Corporation, USA). Muropeptides were separated using organic buffers as described previously.

**Protein expression and purification**

For gene overexpression, *E. coli* Rosetta 2 (DE3) pet22b_bsrP overnight culture was diluted into fresh LB and grown until $OD_{600}$=0.6-0.7 at 37°C. Expression was induced with 0.5 mM IPTG (isopropyl β-D-1-thiogalactopyranoside) for 3 hours at 30°C. Cell were pelleted by centrifugation and resuspended in 50 mM Tris HCl pH 7.2, 150 mM NaCl, 10% glycerol, and Complete Protease Inhibitor Cocktail Tablets (Roche), and lysed with 3 passes through a French press. Clear lysates (30 min, 15,000 rpm, 4°C) were used to purify protein on Ni-NTA agarose column (Qiagen). Protein was eluted with a discontinuous imidazol gradient and visualized by SDS-PAGE electrophoretic protein separation [2].

**Trimmomatic adapter sequences of single-nucleotide polymorphism analysis:**

>PrefixPE/1

AATGATACGGCGACCACCGAGATCTACACTCTTTCCCTACACGACGCTCTTCCGATCT

>PrefixPE/2

CAAGCAGAAGACGGCATACGAGATCGGTCTCGGCATTCCTGCTGAACCGCTCTTCCGATC

T

>PCR_Primer1

AATGATACGGCGACCACCGAGATCTACACTCTTTCCCTACACGACGCTCTTCCGATCT

>PCR_Primer1_rc

AGATCGGAAGAGCGTCGTGTAGGGAAAGAGTGTAGATCTCGGTGGTCGCCGTATCATT

>PCR_Primer2

CAAGCAGAAGACGGCATACGAGATCGGTCTCGGCATTCCTGCTGAACCGCTCTTCCGATC

T

>PCR_Primer2_rc

AGATCGGAAGAGCGGTTCAGCAGGAATGCCGAGACCGATCTCGTATGCCGTCTTCTGCTT

G

>FlowCell1

TTTTTTTTTTAATGATACGGCGACCACCGAGATCTACAC

>FlowCell2

TTTTTTTTTTCAAGCAGAAGACGGCATACGA

>PrefixPE/1

TACACTCTTTCCCTACACGACGCTCTTCCGATCT

>PrefixPE/2

GTGACTGGAGTTCAGACGTGTGCTCTTCCGATCT

>PE1

TACACTCTTTCCCTACACGACGCTCTTCCGATCT

>PE1_rc

AGATCGGAAGAGCGTCGTGTAGGGAAAGAGTGTA

>PE2

GTGACTGGAGTTCAGACGTGTGCTCTTCCGATCT

>PE2_rc

AGATCGGAAGAGCACACGTCTGAACTCCAGTCAC

>PrefixPE/1

TACACTCTTTCCCTACACGACGCTCTTCCGATCT

>PrefixPE/2

GTGACTGGAGTTCAGACGTGTGCTCTTCCGATCT

**Nodulation assay**

*M. sativa* seeds were washed with ethanol and sterilized with 0.1% $HgCl_2$ for 2 min, followed by 6 wash steps with sterile distilled water. The seeds were placed on 1 % agar plates, and stored upside down overnight at room temperature. The germinated seedlings were then placed individually in long test tubes on the surface of 1.2 % agar slants in nitrogen-free Gibson medium [3]. The seedlings were kept at 24°C with 16 hours light and 8 hours dark periods. Two days later the roots (20 plants per condition) were inoculated with 0.2 ml of a *S. medicae* prepared as follows. Starter culture of bacteria was grown overnight in TA liquid medium [4] then bacteria suspension was diluted to $OD_{600}$ = 0.1 and grown further in TA medium in the presence or absence of 20 mM DL-CAN for 12 hours. Subsequently the bacterial cells collected by centrifugation were washed and re-suspended in Gibson liquid medium with or without DL-CAN 20 mM for inoculation of the seedlings. Bacterial dilutions were plated to calculate number of cells per milliliter, and 200 µl of cultures were used for roots inoculation.

The nodulation phenotype of *M. sativa* (nodule number, color and size and general plant phenotype) was scored by visual evaluation from 5 to 13 days post inoculation (dpi) daily and 16 and 23 dpi. Nodules were embedded in 4% agarose and 80 µm nodule sections obtained with vibrotome (MICROM GMBH Germany, model HM650V) sectioning were observed by microscopy (LEICA DMLB2) and pictures were taken and analyzed by QIMAGING/MICROPUBLISHER 3.3RTV camera and using Q-CAPTURE PRO 7 software.

**Supplementary Table 1.** Strains and plasmids used in this study.

| Strains | Media | Temp (°C) | Source/Ref |
|---|---|---|---|
| *Pseudomonas putida* KT2440 | LB | 30 | Courtesy of V. Shingler |
| *Pseudomonas putida* Δ*bsrP* | LB | 30 | This study |
| *Agrobacterium tumefaciens* C58 | LB | 30 | Courtesy of P. Brown |
| *Agrobacterium tumefaciens* C58 *pbp3a*$^{K537R}$ | LB | 30 | This work |
| *Sinorhizobium meliloti* 1021 | LB | 30 | Courtesy of P. Brown |
| *Ensifer arboris* DSM 13375 | LB | 30 | DSMZ collection |
| *Chromobacterium violaceum* CV026 | LB | 30 | CECT collection |
| *Ralstonia pickettii* DSM 6297 | LB | 30 | DSMZ collection |
| *Cupriavidus necator* JNP289 | LB | 30 | Courtesy of E. Diaz |
| *Achromobacter denitrificans* DSM 30026 | LB | 30 | DSMZ collection |
| *Janthinobacterium lividum* CECT 946 T | LB | 30 | CECT collection |
| *Burkholderia gladioli* DSM4285 | LB | 30 | DSMZ collection |
| *Burkholderia cepacia* ATCC 25416 | LB | 30 | Courtesy of L. Thomashow |
| *Xanthomonas campestris* DSM 3586 | LB | 26 | DSMZ collection |
| *Stenotrophomonas rhizophila* DSM 14405 | LB | 30 | DSMZ collection |
| *Buttiauxella agrestis* DSM 4586 | LB | 30 | DSMZ collection |
| *Klebsiella variicola* DSM 15968 | LB | 30 | DSMZ collection |
| *Citrobacter freundii* DSM 30039 | LB | 30 | DSMZ collection |
| *Rahnella aquatilis* DSM 4594 | LB | 30 | DSMZ collection |
| *Pectobacterium carotovorum* DSM 30168 | LB | 26 | DSMZ collection |
| *Pantoea dispersa* DSM 30073 | LB | 30 | DSMZ collection |
| *Pseudomonas syringae* DSM 6693 | LB | 26 | DSMZ collection |
| *Pseudomonas aureofaciens* 30–84 Phz | LB | 30 | Courtesy of L. Thomashow |
| *Escherichia coli* DH5α | LB | 37 | *E. coli* genetic stock |
| *Escherichia coli* DH5α λPIR | LB | 37 | *E. coli* genetic stock |
| *Escherichia coli* Sm10λPIR | LB | 37 | *E. coli* genetic stock |
| *Escherichia coli* Rosetta 2 | LB | 37 | *E. coli* genetic stock |
| *Escherichia coli* S17-1 λPIR | LB | 37 | *E. coli* genetic stock |
| *Escherichia coli* K12 MG1655 | LB | 37 | *E. coli* genetic stock |
| *Sinorhizobium medicae* WSM419 | TA | 30 | Courtesy of E. Kondorosi |

**Supplementary Table 2.** Primers used in this study.

| Name | Description | Sequence (5' - 3') |
|---|---|---|
| FCP1097 | PP3722 NdeI fv | AAAACATATGCCCTTTCGCCGTACC |
| FCP1098 | PP3722 NotI rv | AAAAGCGGCCGCGTCGACGAGTAT |
| FCP1145 | PP3722_P1_XbaI | AAAATCTAGATCATCAGCAGCGACAT |
| FCP1092 | PP3722_P2 | CAATGGCAATTGGTGATTACTCGTGTTC |
| FCP1093 | PP3722_P3 | GAGTAATCACCAATTGCCATTGAAAGGAG |
| FCP1146 | PP3722_P4_XbaI | AAAATCTAGAGCGACGTCACGC |

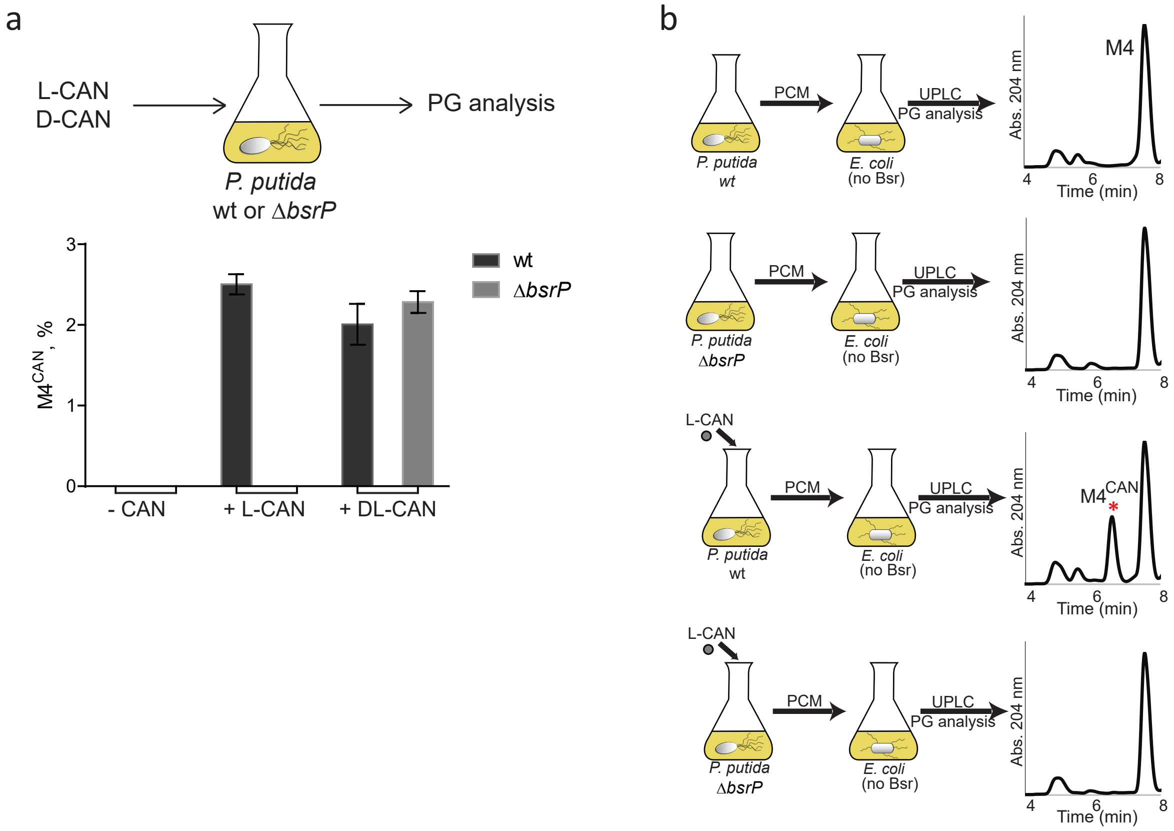


**Fig. S1.** (a) $M4^{CAN}$ accumulation in *P. putida* wild-type and Δ*bsrP* grown without canavanine or supplemented with L-CAN or DL-CAN (5 mM). (b) Cell wall analysis of *E. coli* grown in PCM of *P. putida* wild-type and Δ*bsrP*, which was cultured without or with 5 mM L-canavanine.

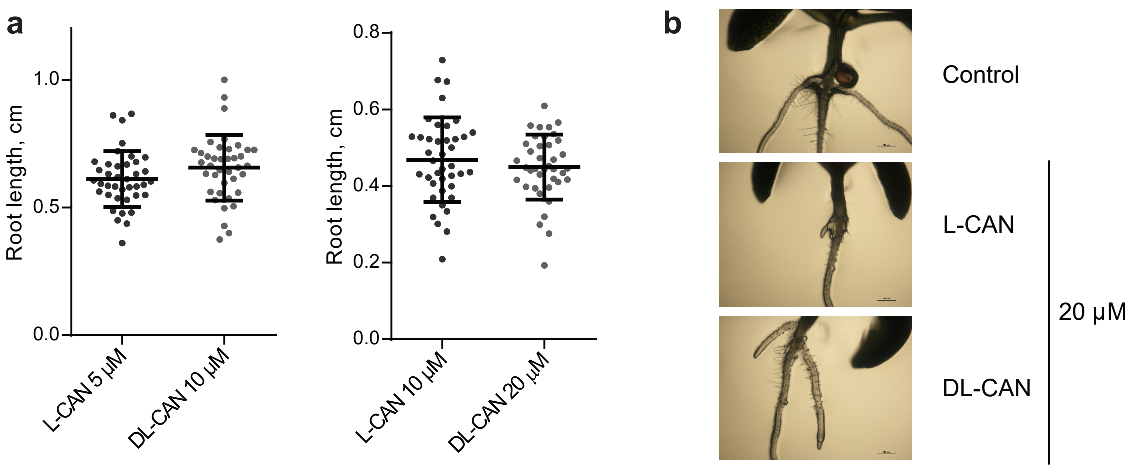


**Fig. S2.** (a )Root length in *A. thaliana* grown on ½ Murashige-Skoog agar supplemented with L-canavanine 5 and 10 µM, and DL-canavanine 10 and 20 µM. (b) Representative pictures of root system in *A. thaliana* grown on ½ Murashige-Skoog agar supplemented with L- or DL-canavanine 20 µM or not (control).

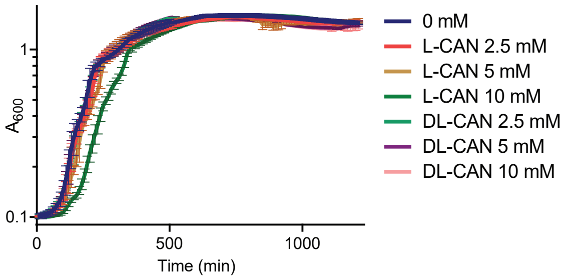


**Fig. S3.** Growth curves of *P. putida* in LB in the absence (0 mM) or presence of L- or DL-canavanine.

a

| Muropeptide | Molecular mass [M+H]$^{+}$ Theoretical | Experimental | Difference |
|---|---|---|---|
| M4$^{CAN}$ | 1029.499 | 1029.526 | 0.027 |
| M5$^{CAN}$ | 1100.536 | 1100.579 | 0.043 |
| D34$^{CAN}$ | 1881.859 | 1881.879 | 0.02 |
| D44$^{CAN}$ | 1952.896 | 1952.986 | 0.02 |
| D45$^{CAN}$ | 2023.933 | 2024.016 | 0.083 |

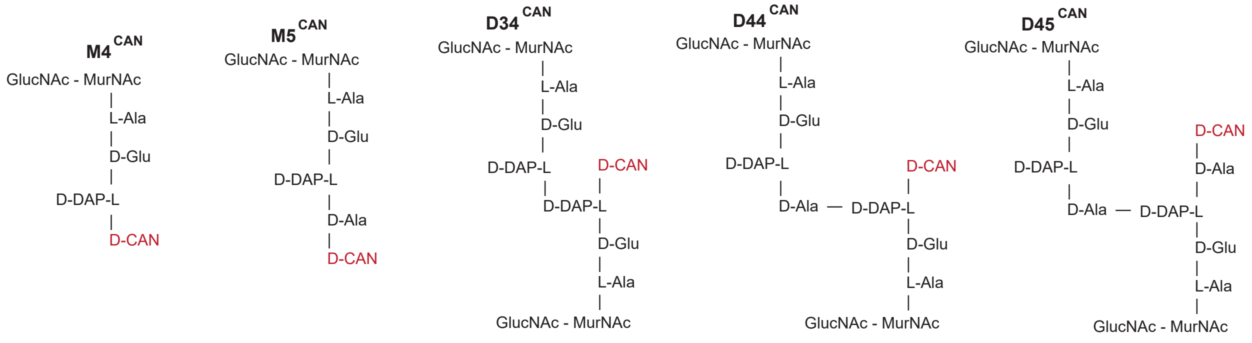


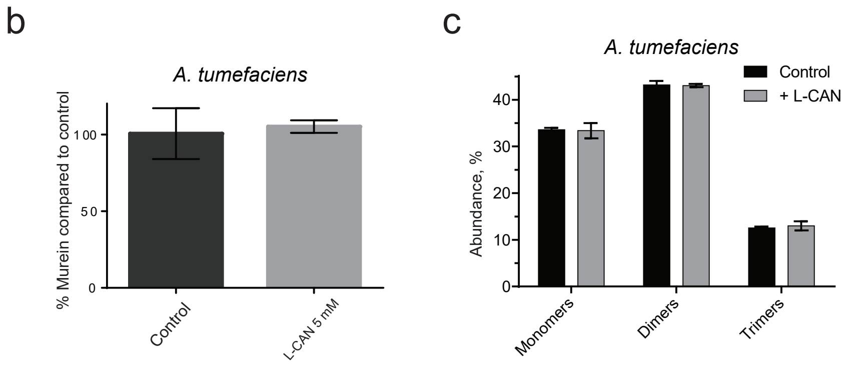


**Fig. S4.** (a) Masses and structure of CAN-modified muropeptides. (b) Peptidoglycan amount quantification in 5 mM L-canavanine supplemented *A. tumefaciens* cultures normalized to control (no canavanine). (c) Abundance of monomers, dimers and trimers in *A. tumefaciens*, grown in LB medium (control) or LB medium supplemented with L-canavanine 5 mM.

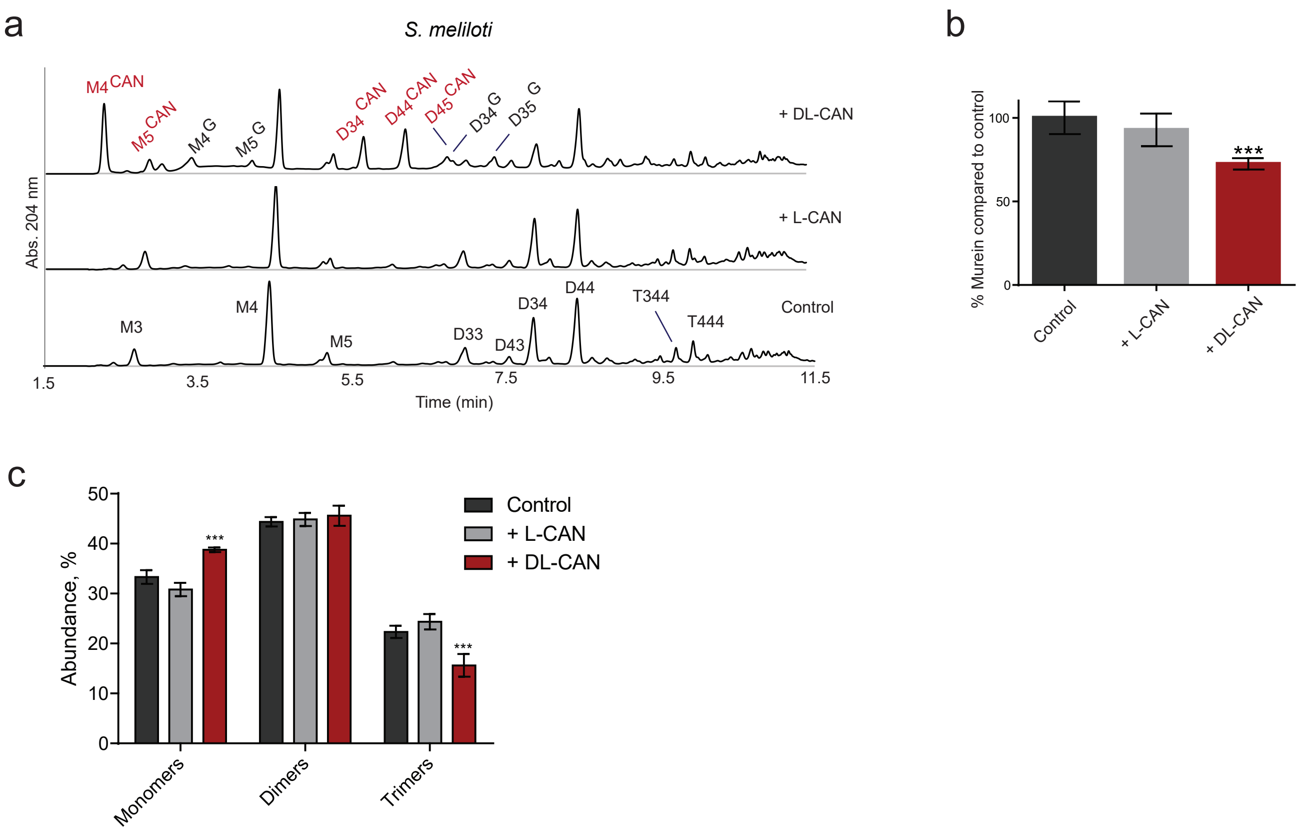


**Fig. S5.** (a) Representative peptidoglycan profiles of *S. meliloti* supplemented with L- or DL-canavanine 1.5 mM or not supplemented (control). (b) Peptidoglycan amount quantification in 1.5 mM L-canavanine and 1.5 mM DL-canavanine supplemented *S. meliloti* cultures normalized to control (no canavanine). P value < 0.0001 (***). (c) Abundance of monomers, dimers and trimers in *S. meliloti*, grown in LB medium (control) or LB medium supplemented with L- or DL-canavanine 1.5 mM. P value < 0.0001 (***).

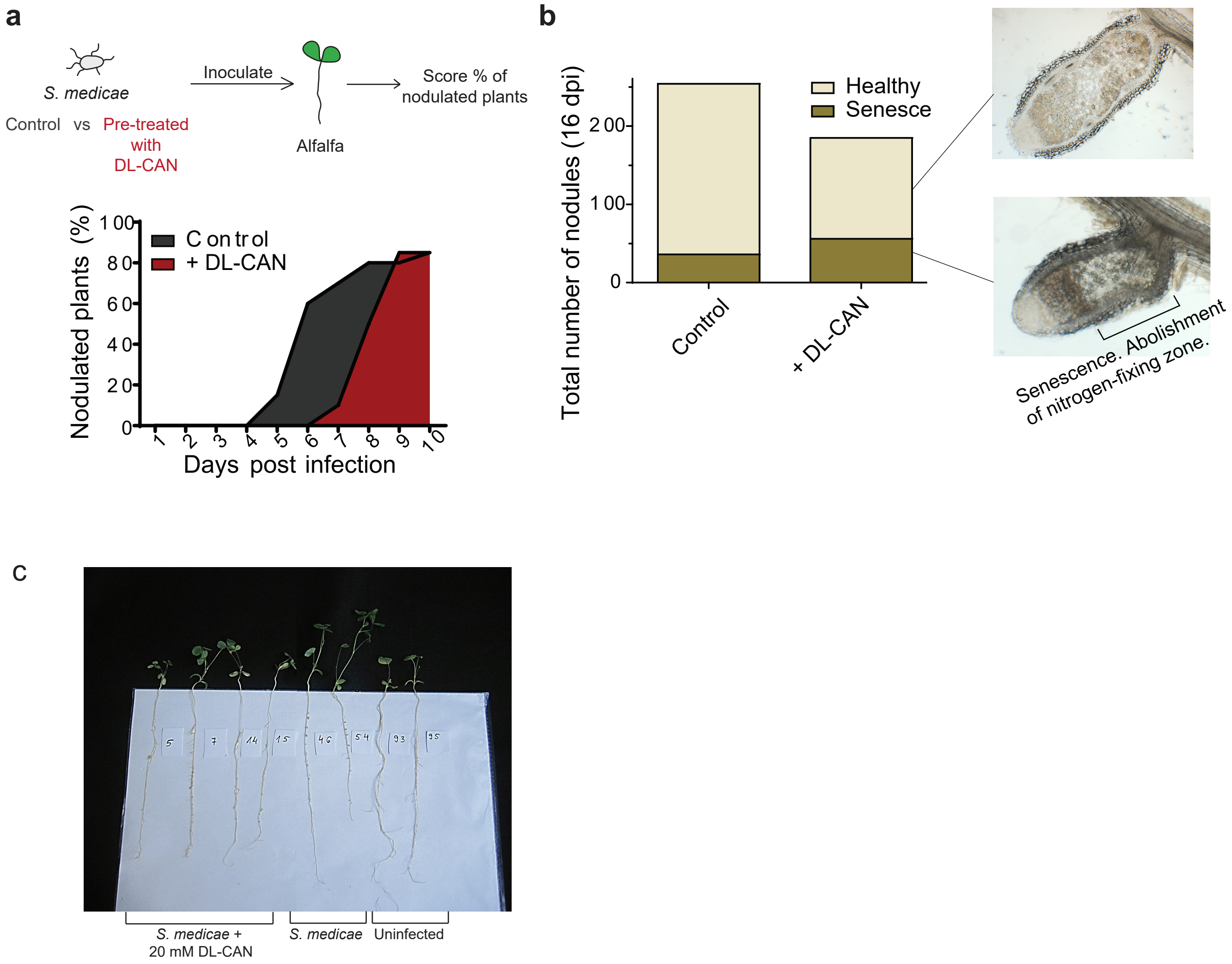


**Fig. S6.** (a) Scheme of the experiment and nodulation kinetics of *Medicago sativa*, infected with *S. medicae* WSM419 pre-incubated with DL-canavanine 20 mM or not (control). (b) Total nodule count and amount of healthy and senescent nodules in *Medicago sativa* (20 plants per condition) 16 days post infection, infected with *S. medicae* WSM419 pre-incubated with DL-canavanine 20 mM or not (control). Pictures show representative cross-sections of healthy and senescent nodules. (c) Picture of *Medicago sativa* representative plants, inoculated with *S. medicae* pre-treated with DL-canavanine 20 mM, non-pretreated, and uninfected *Medicago sativa* plants.

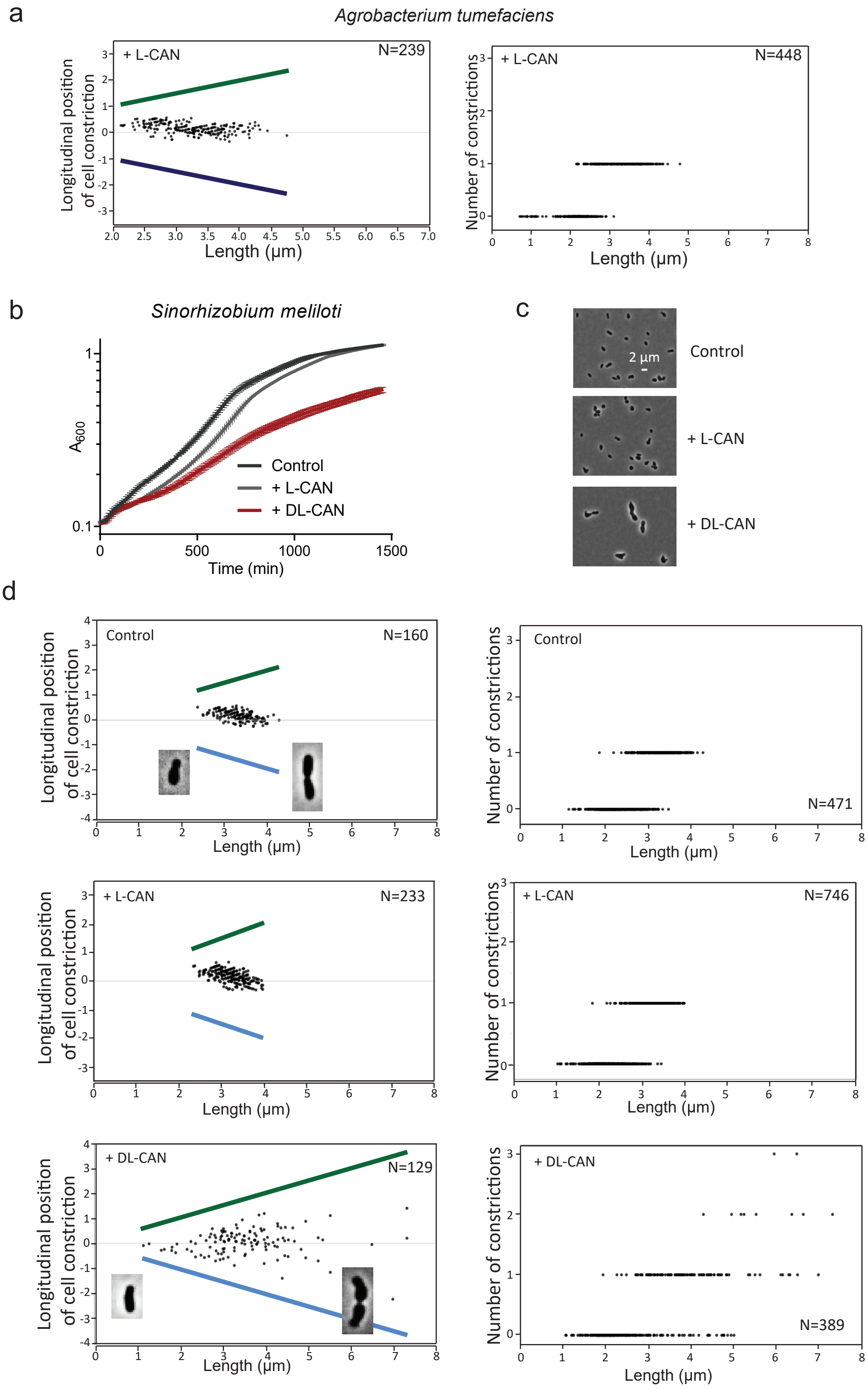


**Fig. S7.** (a) Longitudinal position of cell constriction and number of constrictions in *A. tumefaciens* cells with L-canavanine 10 mM. (b) Growth curves of *S. meliloti* in LB medium supplemented with L- or DL-canavanine 2.5 mM or not (0 mM). (c) Phase contrast images of *S. meliloti* cells without (control) or supplemented with L- or DL-canavanine 3 mM. Scale bar 2 µm. (d) Longitudinal position of cell constriction and number of constrictions in *S. meliloti* cells with L- or DL-canavanine 3 mM or without addition of canavanine (control).

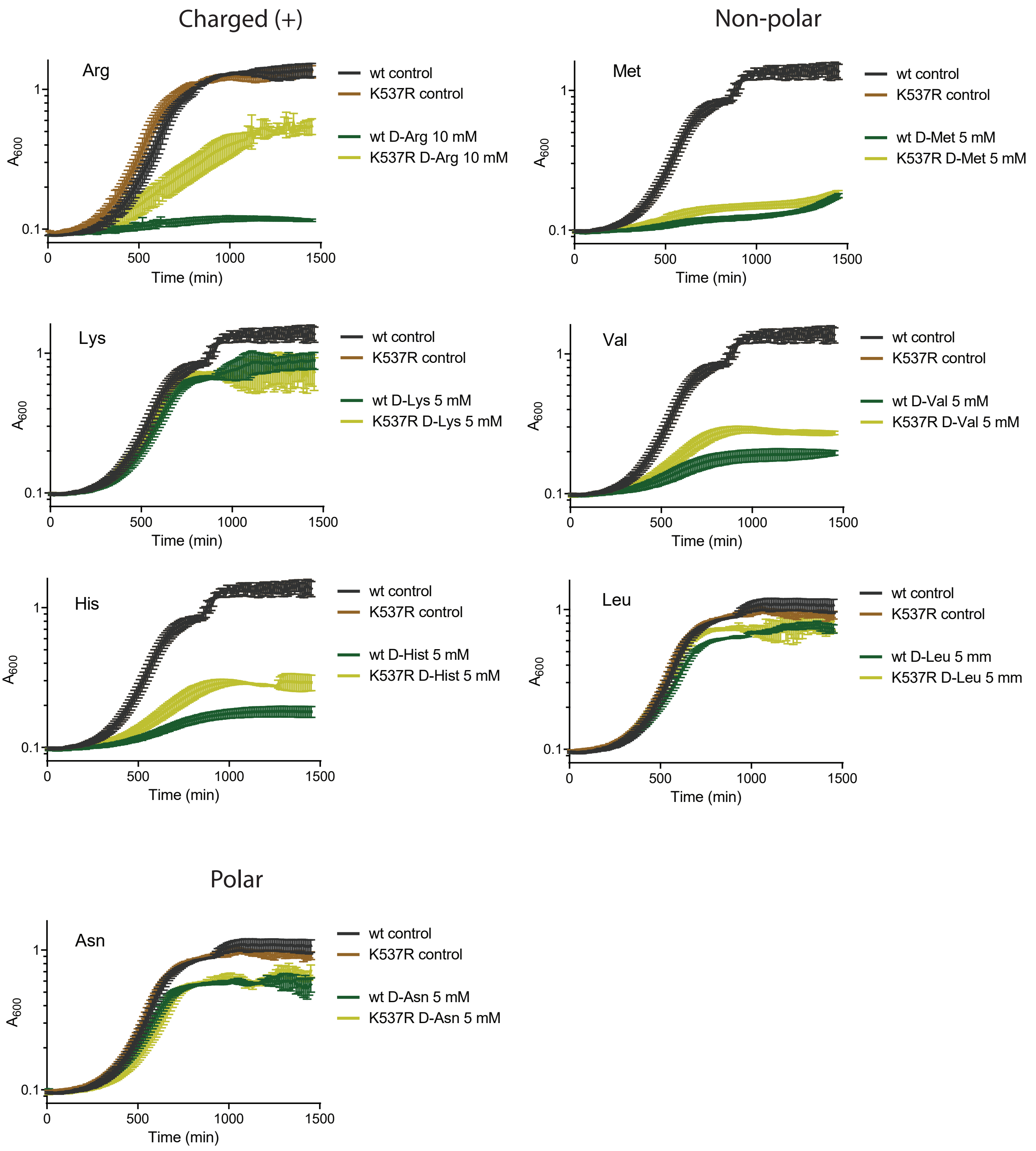


**Fig. S8.** Growth curves of *A. tumefaciens* wild-type and PBP3a$^{K537R}$ without and with supplementation of different D-amino acids.

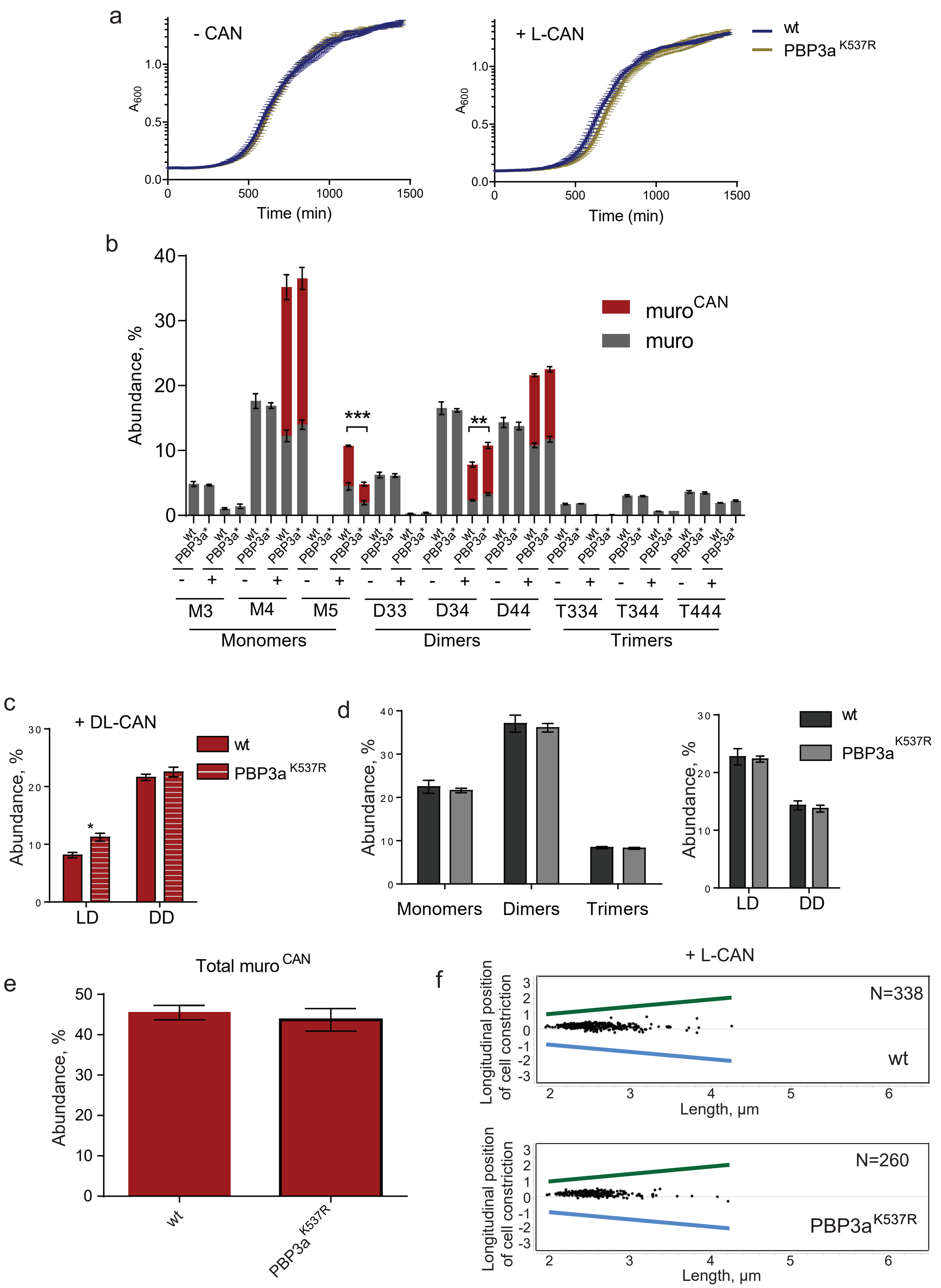


**Fig. S9.** (a) Growth curves of *A. tumefaciens* wild-type and PBP3a$^{K537R}$ in LB medium or in LB medium supplemented by L-canavanine 10 mM. (b) Abundance of D-canavanine-containing muropeptides in *A. tumefaciens* wild-type and PBP3a$^{K537R}$ grown with or without DL-canavanine 10 mM. P-value < 0.005 (**) and < 0.0001 (***). (c) Abundance of LD- and DD-crosslinked muropeptides in *A. tumefaciens* wild-type and PBP3a$^{K537R}$ supplemented with 10 mM DL-canavanine. (d) Abundance of monomers, dimers and trimers in *A. tumefaciens* wild-type and PBP3a$^{K537R}$, grown in LB medium (control). (e) Abundance of D-canavanine-containing muropeptides in *A. tumefaciens* wild-type and PBP3a$^{K537R}$ supplemented with 10 mM DL-canavanine. (f) Longitudinal position of cell constriction in *A. tumefaciens* wild-type and PBP3a$^{K537R}$ cells with L-canavanine 7.5 mM.
